# Description of *Phylloneura rupestris* sp. nov. (Odonata, Platycnemididae) from the Western Ghats, India with notes on its reproductive behavior

**DOI:** 10.1101/2023.09.13.557666

**Authors:** Ayikkara Vivek Chandran, Reji Chandran, S R Suraj, Subin K. Jose, Pankaj Koparde

## Abstract

*Phylloneura* Fraser, 1922 is a genus of damselflies which was till date regarded monotypic. It is represented by the nominate species, *Phylloneura westermanni* (Hagen in Selys, 1860) which is endemic to the Western Ghats in India. In our study, we came across a population of *Phylloneura* near Ponmudi hills, Thiruvananthapuram, which displayed morphological and behavioral differences from *P. westermanni*. We describe it as a species new to science with detailed photographs and illustrations. We also provide notes on its reproductive behavior.

## Introduction

*Phylloneura westermanni* (Hagen in Selys, 1860) is a species of damselfly belonging to Family Platycnemididae (Jacobson and Bianchi, 1905). It is seen in colonies, associated with Myristica swamps and associated streams, and hence has the common name Myristica Bambootail (Subramanian 2009). Till date, it has remained the sole described species of the genus and is considered to be Near Threatened by IUCN (Subramanian 2011). It is endemic to the Western Ghats of India and has been recorded only between the Nilgiri Hills and Sharavathi Valley, north of the Palghat Gap (Subramanian et al. 2018). The body of *P. westermanni* is jet black marked with azure blue and it is distinguishable from other bambootails of the Western Ghats by the following features: segment 7 of the abdomen with a broad blue apical ring, greater number of postnodal nervures (28-31 in forewings and 26-27 in hindwings) and presence of many double cells between main nervures (Fraser 1933).

As part of our ongoing study on the odonate fauna of the Western Ghats, we came across a population of *Phylloneura* near Ponmudi hills, a part of the Agasthyamalai hill range, that differed in markings on the 7^th^ abdominal segment. This population also showed peculiar breeding behavior, we observed multiple pairs ovipositing on moss growing in the rills of a seasonal stream flowing steeply over a rocky cliff. We collected some individuals of this population and compared their morphology with that of *P. westermanni* specimens collected from Wayanad, which allows us to describe the population from Ponmudi as a new species.

## Material and methods

Field observations of the *Phylloneura* population of Merchiston estate, Ponmudi (8.749690° N, 77.127098° E, 815 m above MSL) were made using Nikon 8x42 Aculon A211 binoculars. Reproductive activities of a pair were timed using a stopwatch. Photographs of live insects in the field were taken using a Nikon Z6II mirrorless camera, Nikon 200-500 mm telephoto lens and Nikkor Z MC 105 mm macro lens. Three male and a female *Phylloneura* specimens from Merchiston estate, Ponmudi were collected using a butterfly net. One male specimen was preserved in absolute ethanol and the rest were preserved dry. For comparison, two male *Phylloneura westermanni* specimens were caught from Thirunelli, Wayanad (11.913150° N, 75.971120° E, 848 m above MSL) using a butterfly net, one of which was preserved in absolute ethanol and the other was preserved dry (Figure 1). Morphological data for the specimens were made using a stereomicroscope (SkiHi TDLED-1005, India), and they were photographed using a mirror-less digital camera (Sony a7III body, Sony 90mm macro lens and Raynox DCR-250 super macro lens). The male specimens preserved in ethanol were dissected to study their genitalia. After studying the genital ligulae under microscope, they were preserved in vials filled with absolute ethanol. Diagrams of genital ligulae and caudal appendages were made by tracing their photographs using the android application Sketchbook. Descriptive terminology follows Garrison et al. (2010). All measurements were taken using a digital Vernier calliper (ZHART CT-ZT-VERNIER).

**Figure 1:**
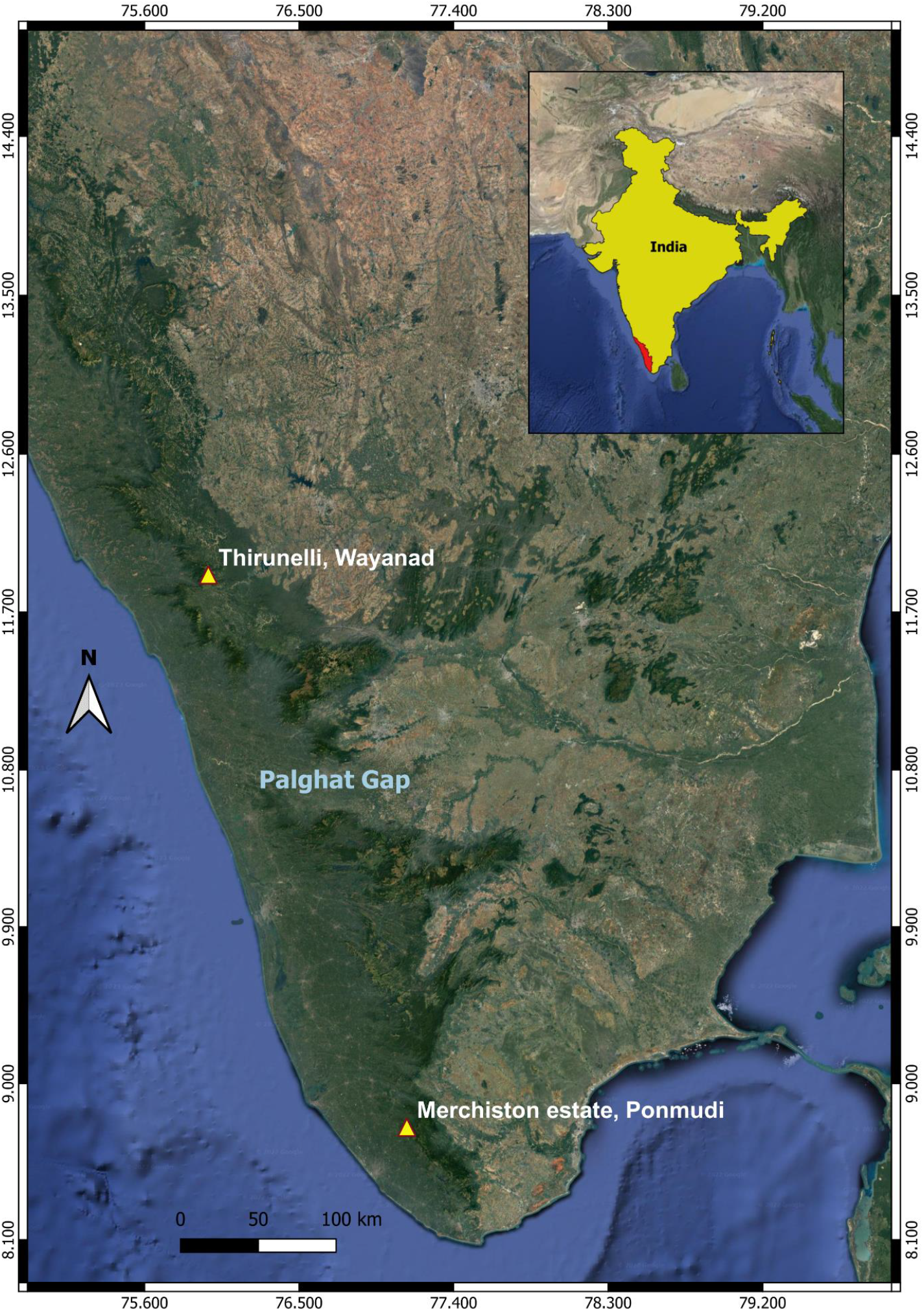
Map of Southern Western Ghats showing locations of collection of *Phylloneura rupestris* sp. nov. (Merchiston estate, Ponmudi) and *P. westermanni* (Thirunelli, Wayanad)

Type specimens and preserved genital ligulae are now in the insect collections of the Department of Geology and Environmental Science, Christ College (Autonomous), Irinjalakuda (ICDGE). The holotype will be deposited in the National Designated Repository for Fauna, Western Ghat Regional Centre, Zoological Survey of India (WGRC, ZSI) in due course. The following abbreviations are used in this study: FW= Fore wing, HW= Hind wing, Ab= Anal bridge, Ax= Antenodal nervures, Px= Postnodal nervures, Pt= Pterostigma, S1-S10= first to tenth abdominal segments. Specimen label data are quoted verbatim.

### Systematic accounts

Order Odonata Fabricius, 1793

Suborder Zygoptera Selys, 1854

Superfamily Coenagrionoidea Kirby, 1890

Family Platycnemididae Jacobson and Bianchi, 1905

**Genus Phylloneura** Fraser, 1922

Type species: *Phylloneura westermanni* (Hagen in Selys, 1860)

***Phylloneura rupestris*** Chandran, Chandran and Jose sp. nov. urn:lsid:zoobank.org:pub:86F47549-9E5F-407B-8DA3-B639DE2C8CC6

#### Type material

Holotype. ♂, labelled as “Holotype *Phylloneura rupestris* Chandran, Chandran, Jose, India, Kerala, Thiruvananthapuram, Ponmudi [8.749690° N, 77.127098° E], elevation 850 m above MSL, 28 vi 2023, Coll. A. Vivek Chandran” (WGRC, ZSI).

Paratypes. 2 ♂ and 1 ♀, data as holotype (ICDGE).

#### Diagnosis

*Phylloneura rupestris* sp. nov. can be distinguished in the field from its only congener, *P. westermanni*, by the broad azure blue marking on S7 that extends to 3/4^th^ of the segment dorsolaterally (S7 has only an azure blue apical ring occupying 1/5^th^ of the segment in *P. westermanni*) (Figure 2). The following features become evident upon close inspection of specimens and in combination, help to distinguish *P. rupestris* sp. nov. from *P. westermanni* (Figures 3 and 4): Pt rectangular with a length of 1.5 mm (Pt squarish with a length of 1 mm in *P. westermanni*). Cerci thicker with broad ventral processes (cerci thinner with narrower ventral processes in *P. westermanni*). Tip of paraprocts rounded (Tip of paraprocts tapers to an acute point in *P. westermanni*). In the apical part of male genital ligula, the two outer processes are hammer shaped and the two inner processes are thicker (tips of the two outer processes end as small lobes and the two inner processes are thinner in *P. westermanni*).

**Figure 2:**
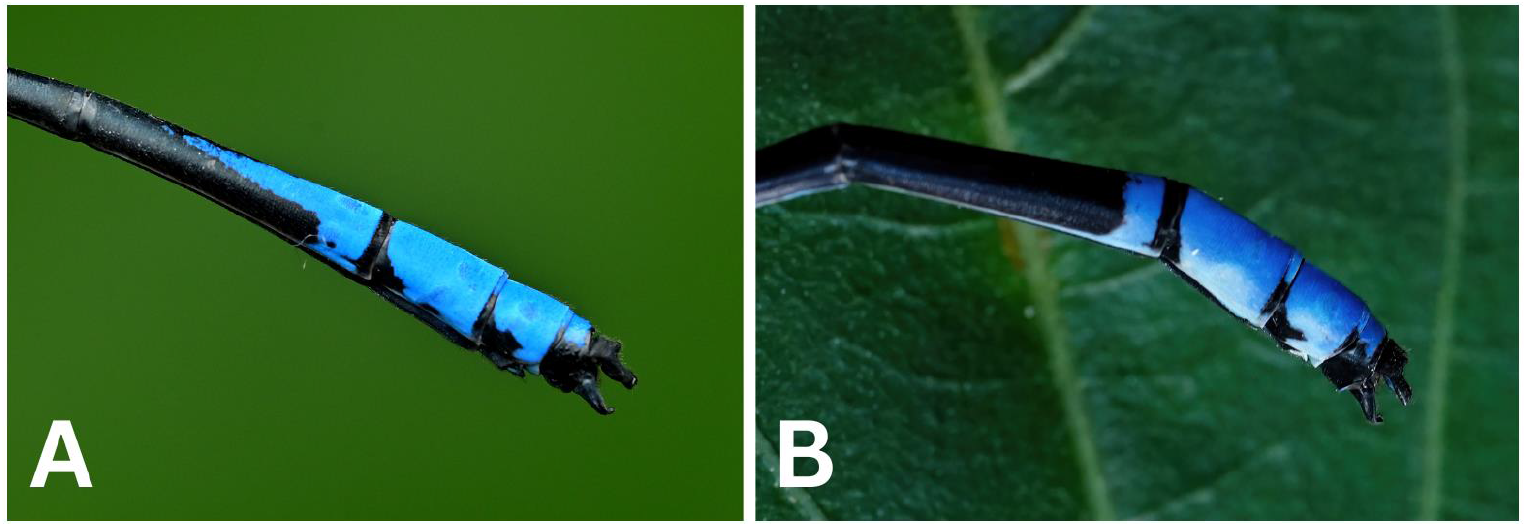
Last three abdominal segments of - A. *Phylloneura rupestris* sp. nov. and B. *P. westermanni* showing the difference in markings on S7.

**Figure 3:**
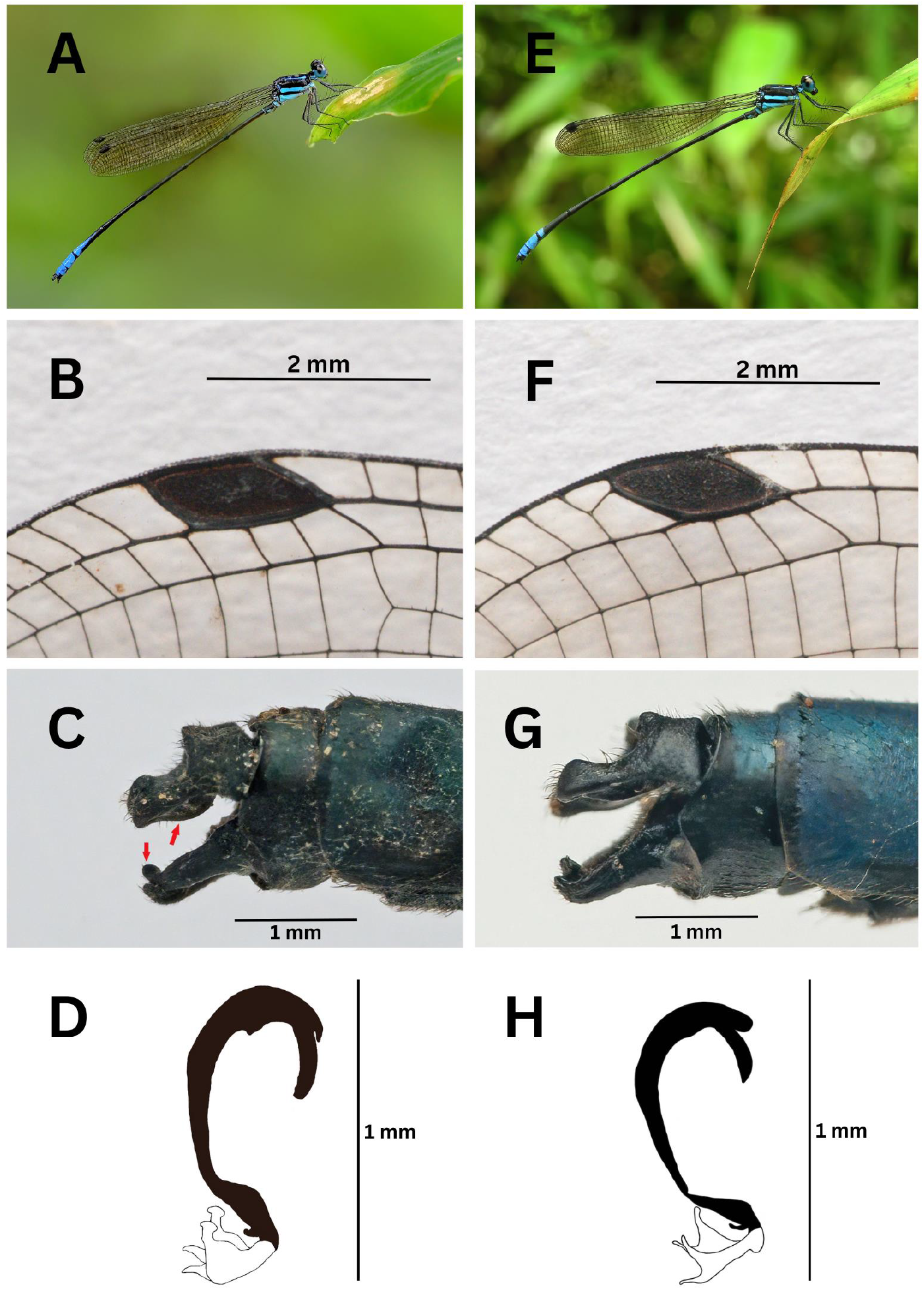
*Phylloneura rupestris* sp. nov.: A- In habitat, B- Pterostigma, C- Caudal appendages, D- Genital ligula; *P. westermanni*: E- In habitat, F- Pterostigma, G- Caudal appendages, H- Genital ligula.

**Figure 4:**
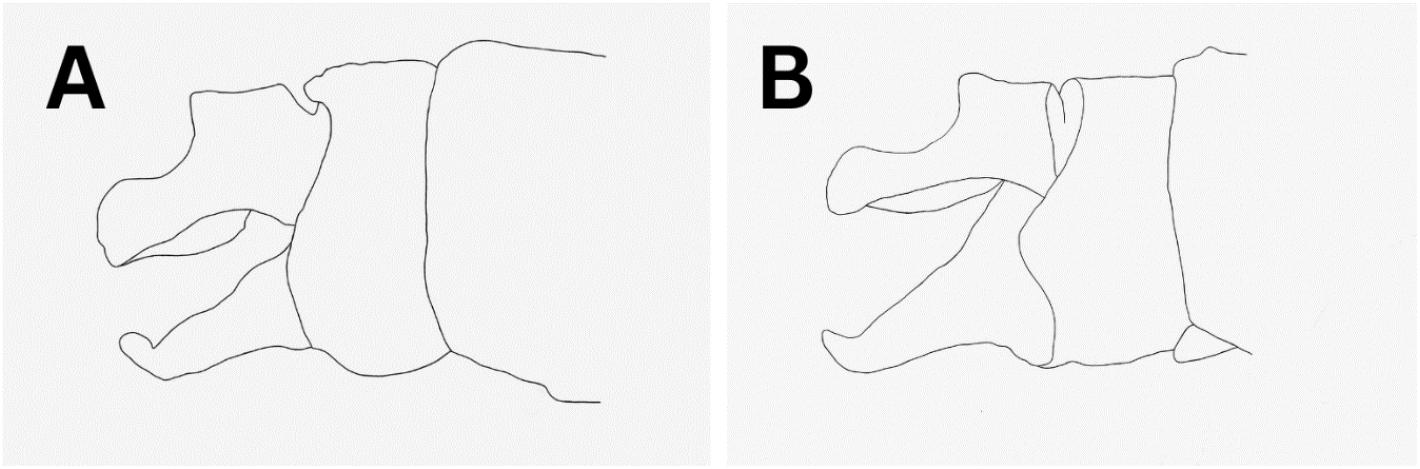
Caudal appendages of : A- *Phylloneura rupestris* sp. nov. and B- *P. westermanni*, traced from original photographs.

It must be noted here that in the description of *P. westermanni* given by Fraser (1922, 1933), the “apical half” of S7 is mentioned to be azure blue. This, however, is not correct, and as evidenced by photographs published later on (Subramanian 2009, Subramanian 2018, Nair et al. 2021, Gopalan et al. 2022, Subramanian et al. 2022) and specimens collected in this study, only the apical 1/5^th^ of S7 is azure blue that appears as a thick ‘ring’.

#### Description of holotype

(Figure 5). Male. Total length 60 mm; abdomen 49 mm; HW 36 mm.

**Figure 5:**
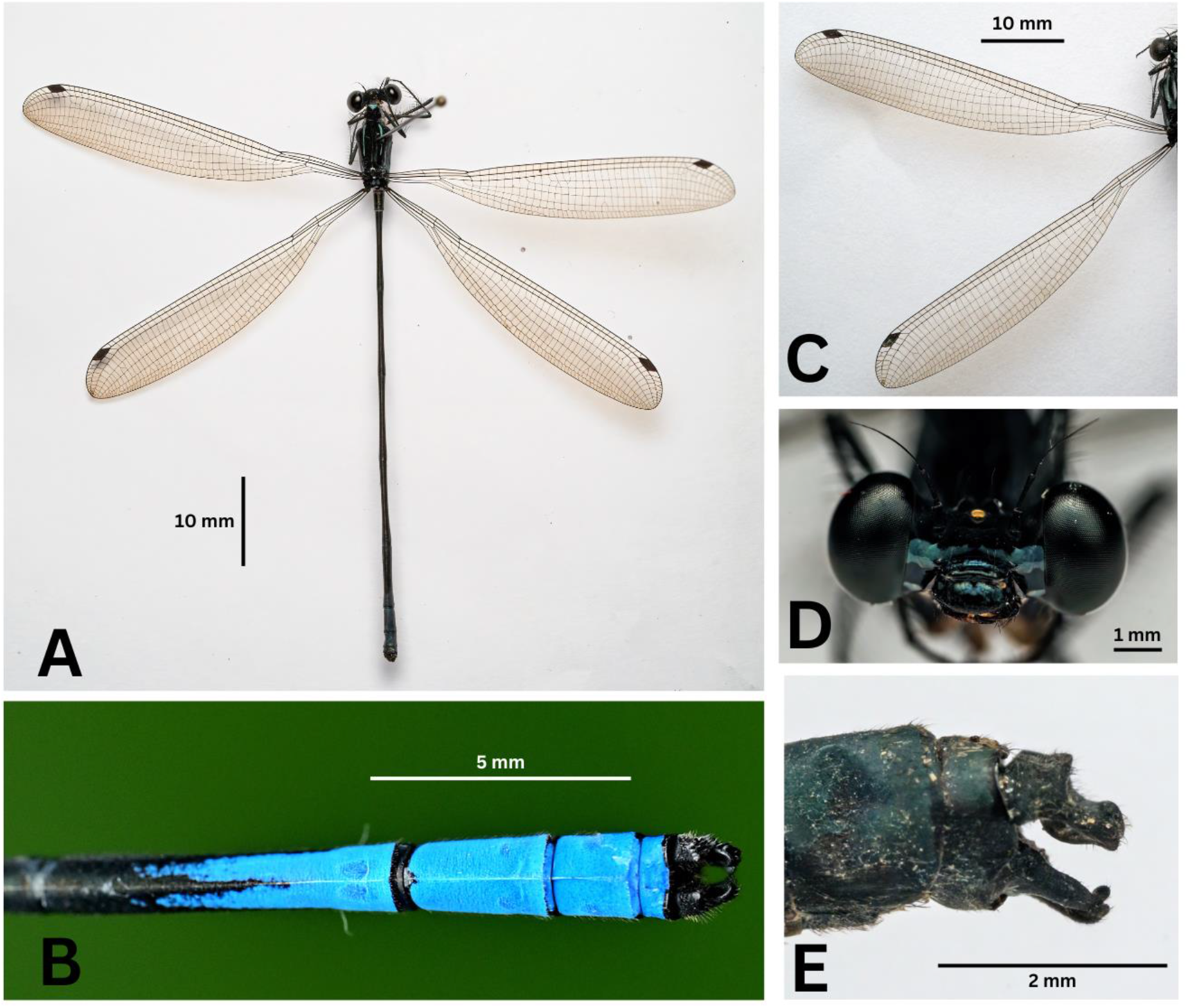
*Phylloneura rupestris* sp. nov. holotype male - A. Habitus, B- Last three abdominal segments in dorsal view, C- Wings, D- Head in frontal view, E- Caudal appendages in lateral view.

Head. Middle and lateral lobes of labium white, tipped black; labrum, anteclypeus and postclypeus steely blue-black, outer margin of postclypeus produced into short horns on both ends; bases of mandibles black, with a central blue spot; genae blue, with a black band which is confluent with the black of clypeus; antefrons black, but the blue of genae spreads onto it narrowly; postfrons fully black; rest of the head black, including the undersurface; lower 2/3^rd^ of eyes ultramarine blue, capped with black; ocelli translucent white.

##### Prothorax

Posterior lobe fully black; middle lobe almost fully black dorsally, but with a short ingress of azure blue from anterior lobe, azure blue laterally; propleuron azure blue; anterior lobe almost fully azure blue, but with short ingress of black from middle lobe at two points dorsally.

##### Pterothorax

Jet black marked with azure blue as follows-middorsal carina and mesepisternum black; very narrow antehumeral blue stripes; mesepimeron black with a white spot at the anterior end on each side; metepisternum black with a broad blue stripe; mesinfraepisternum black, with only its lower end blue; metepimeron blue; underside of thorax bluish white.

##### Legs

Black, coxae and basal half of trochanters blue; extensor surface of tibiae pale yellow.

##### Wings

Hyaline, enfumed lightly; Ab complete, arched at origin, inclining towards hinder border of wing distally; Pt dark reddish brown, rectangular, 1.5 mm in length, covering 3 cells, braced between thick, black nervures; Ax 2 in all wings; Px 33 in left FW, 32 in right FW, 29 in both HW; many double cells in all wings, mostly towards the tips of FW.

##### Abdomen

Black, marked with blue as follows-S1 broadly blue, with black basal and apical margins, the former produced into a median spot dorsally; S2 with a broad blue ventrolateral spot; S3 and S4 have very narrow basal annules; S5 has an almost obsolete basal spot; S6 fully black; S7 with a broad black apical margin from where a large blue stripe extends to 3/4^th^ of the segment towards its base, this stripe split centrally from half its length; S8 and S9 entirely blue on dorsum, except for very narrow basal black annules, laterally almost fully blue, but black irregularly towards the bases and apices; S10 blue on dorsum, black laterally.

##### Caudal appendages

Black, 1.5 times the length of S10, covered irregularly with light brown hairs; cerci broad at base, with an upper obtuse tubercle at base, and another at the tip, a broad ventral process at its apical 3/4^th^ portion; paraprocts broad at base, sloping downwards and tapering gradually, tip rounded and directed upward.

*Paratype* 1. Male. Total length 60 mm; abdomen 49 mm; HW 35 mm.

Px 30 in both FW, 27 in left HW and 26 in right HW; rest agreeing with holotype.

##### Genital ligula

As illustrated in Figure 3D, first segment broad and curved; second segment short and featureless; tip of third segment produced into four filaments, the two outer ones hammer shaped, the two inner ones thick, conical, curved slightly at tips.

*Paratype* 2. Male. Total length 60 mm; abdomen 49 mm; HW 34.5 mm.

Px 31 in left FW, 30 in right FW, 29 in left HW and 26 in right HW; rest agreeing with holotype.

*Paratype* 3. (Figure 6). Female. Total length 58 mm; abdomen 47 mm; HW 34.5 mm.

**Figure 6:**
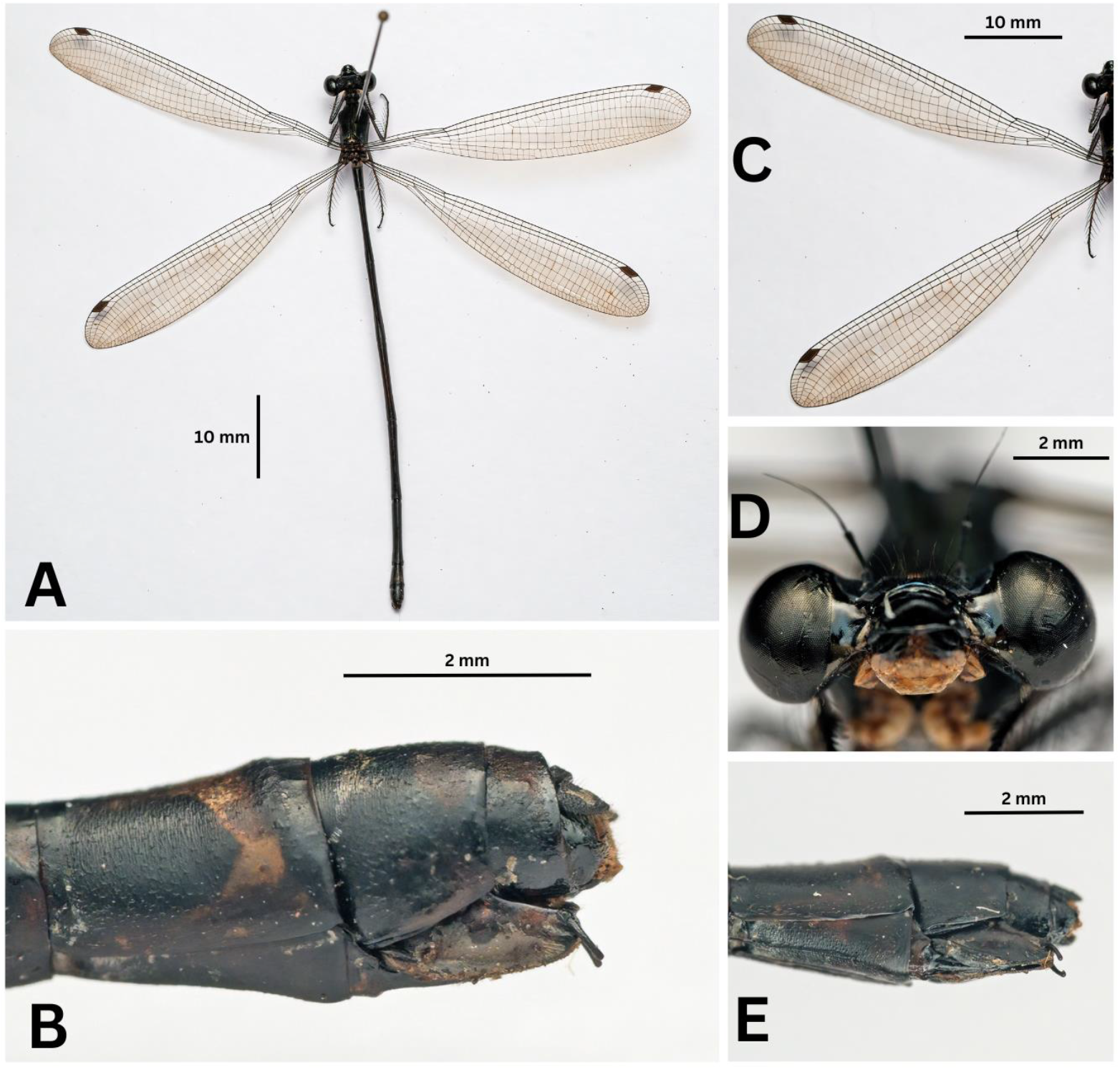
*Phylloneura rupestris* sp. nov. paratype female - A. Habitus, B- Last three abdominal segments in lateral view, C- Wings, D- Head in frontal view, E- Last three abdominal segments in ventrolateral view.

Similar to the male, except for sexual differences and blue markings on the last abdominal segments. Abdomen shorter and more robust; S7 with an apical blue ring, thinner on dorsum, expanding laterally, but not joining ventrally; S8 and S9 with large triangular dorsal blue spots; dorsum of S10 completely blue.

##### Caudal appendages

Black, covered with light brown hairs; cerci short and conical; vulvar scale blue, tipped with black, not reaching the end of abdomen; ovipositor black, robust.

#### Remarks

Male *Phylloneura rupestris* sp. nov. can at once be told apart in the field from male *P. westermanni* by the extensive blue marking on S7, which in the latter is reduced to a thick apical ring. Also, female *P. rupestris* sp. nov. has a distinctive apical blue ring on S7, which in the female *P. westermanni* is obscure. When a male specimen is in hand, a close look at the caudal appendages with a magnifying lens would help in confirming the species, as *P. rupestris* sp. nov. has considerably thicker cerci and rounded tips of paraprocts. Moreover, as per current knowledge, ranges of these two species do not overlap.

#### Etymology

The species epithet *rupestris* in Latin means “that lives on cliffs or rocks” and acknowledges the species’ habitat preference.

#### Distribution

Presently, the species is known only from the type locality (Ponmudi hills, southern Western Ghats), but it could be present in similar habitats (seasonal streams flowing over rocky cliffs at mid elevations) in the Agasthyamalai hills, as these form a continuous chain of forested hills in the southern Western Ghats.

#### Temporal distribution

Recorded in June (30) and July (2).

#### Natural history

A colony of 30 individuals was seen perched on vegetation around a steep rocky cliff over which rills of a seasonal stream flowed (Figure 7). The average depth of the stream, measured using a meter tape was 0.14 m. The sky was overcast and there was light drizzle. Most of the observed individuals were males and females were seen only while mating. They were occasionally making short, slow flights, probably after flying insect prey, and returning to their perches.

**Figure 7:**
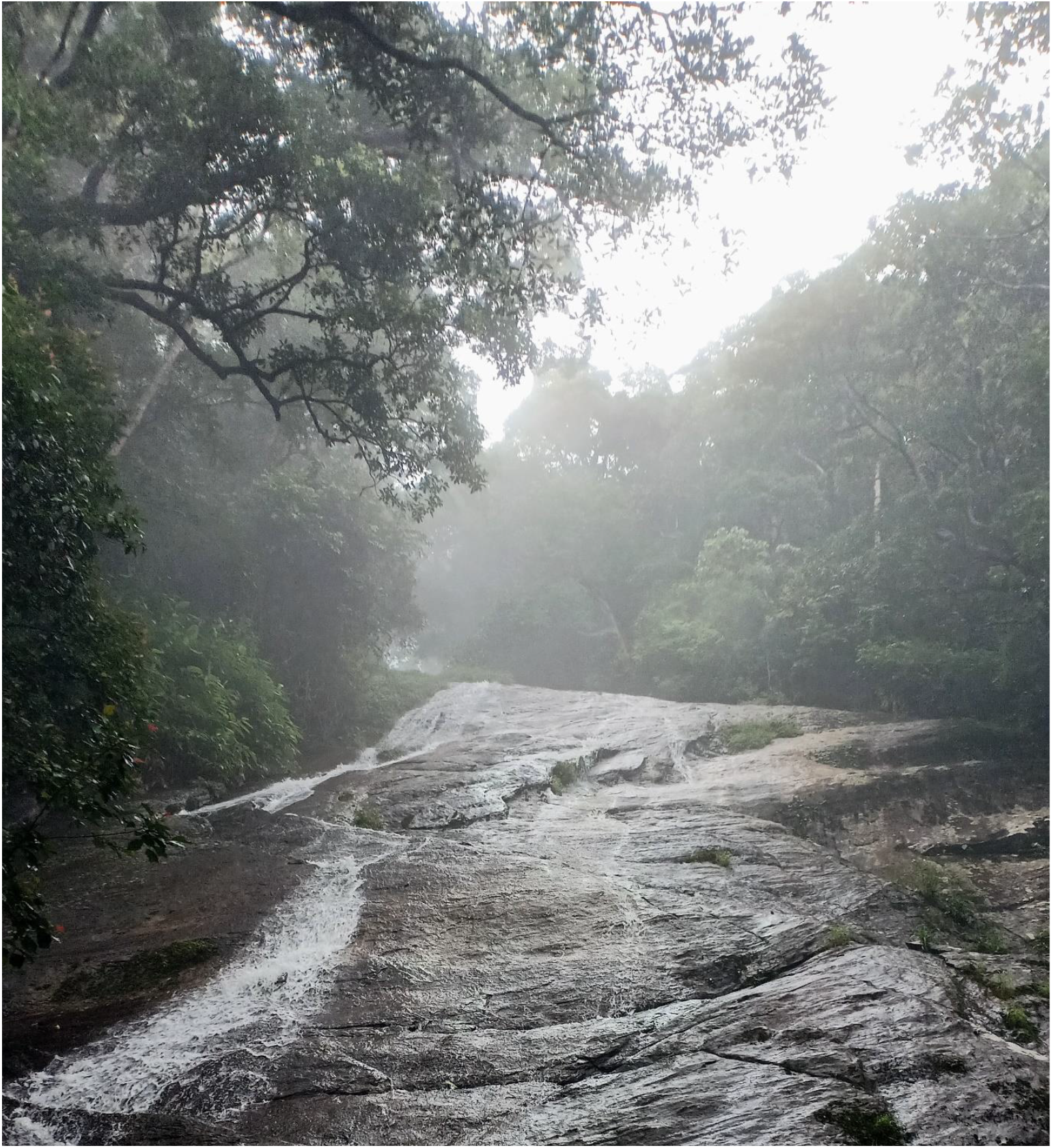
Habitat of *Phylloneura rupestris* sp. nov.

#### Reproductive behavior

Oviposition of 6 pairs were seen, but we could observe, photograph and time the entire chain of reproductive activities of only one pair (Figure 8). The time of each activity in seconds (s) is given in parenthesis-First, the male descended from its perch very slowly, hovered just above the stream and dipped its abdomen in water. Thereafter, it flew back in the direction of its former perch and caught a female in tandem (22 s). The male performed intra-male sperm translocation with the female held in tandem (10 s). Next, the pair flew around in tandem over the stream and slowly descended (18 s). The female oviposited on a submerged mat of moss in the stream, with the male holding it in tandem (10 minutes). The pair took flight again, perched on another patch of moss in the stream about 3 meters away and oviposited again (4 minutes). Thereafter, the male, with the female held hanging in tandem, hung on streamside vegetation (41 s). After this, they separated and flew in different directions. The entire reproductive activity of the observed pair, thus, lasted for 15.51 minutes.

**Figure 8:**
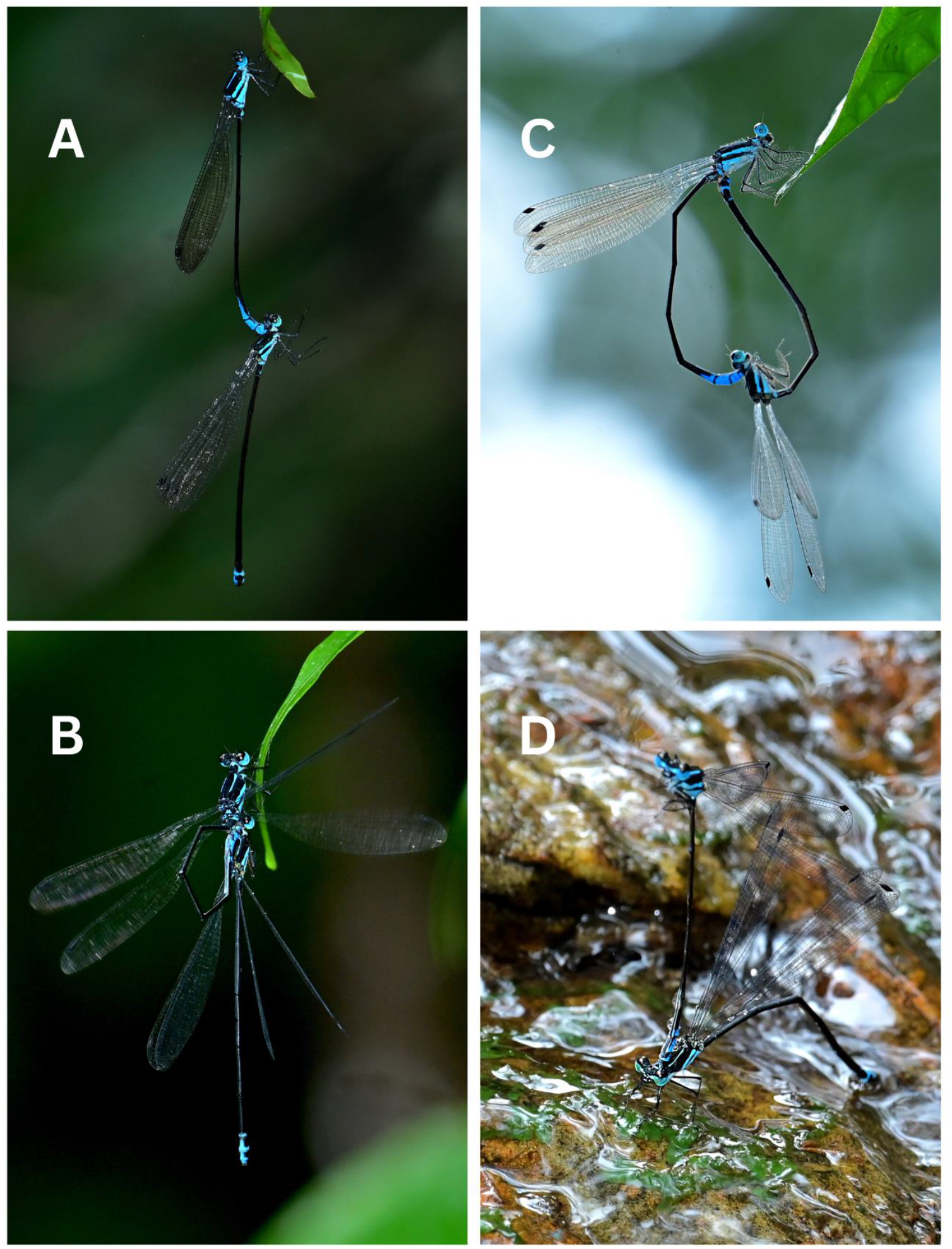
Reproductive behavior of *Phylloneura rupestris* sp. nov. : A- Tandem, B- Intra-male sperm translocation, C- Copula, D- Oviposition.

All the 6 observed pairs oviposited on moss growing in the seasonal rills over the rocky cliff. This ovipositing technique is in sharp contrast to that of *P. westermanni*, whose pairs were observed ovipositing on surface roots of riparian trees (Figure 9).

**Figure 9:**
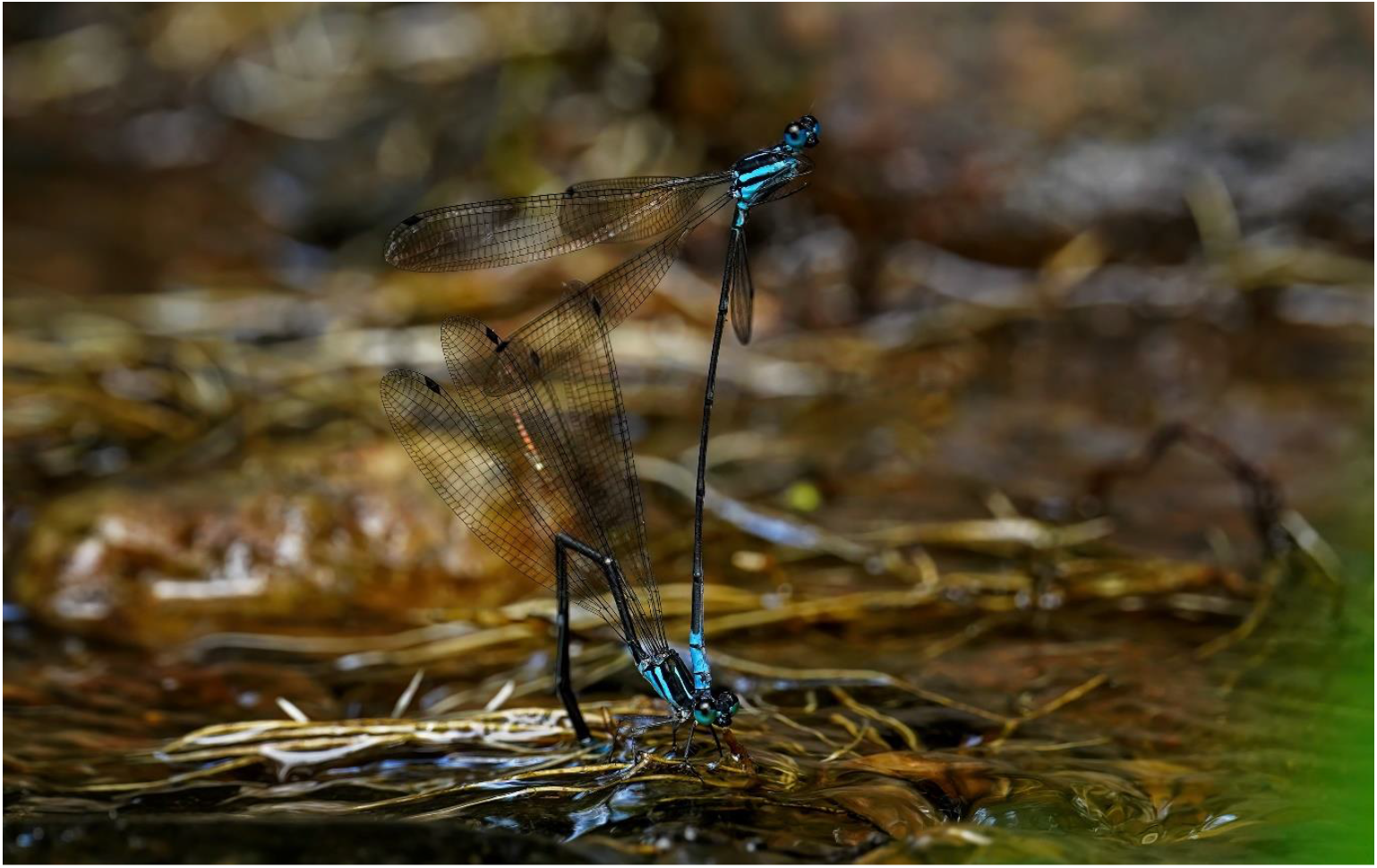
A pair of *P. westermanni* ovipositing on surface roots of a riparian tree.

## Discussion

Biodiversity of insects is threatened worldwide, due to habitat loss, pollution, pathogens and invasive species, and climate change (Sánchez-Bayo and Wyckhuys, 2019). An IUCN assessment indicates that 10% of the world’s Odonata are threatened with extinction, although the study only covered a quarter of all species known (Clausnitzer et al. 2009). The Western Ghats is a global biodiversity hotspot (Myers et al. 2000) with several new species of Odonata described in recent times (Payra et al. 2023; Vijayakumaran et al. 2022; Joshi et al. 2022; Rangnekar et al. 2019). However, *P. rupestris* sp. nov. is the first Platycnemidid to be described from this landscape after 1931. This study suggests that there could be more odonate species, even in groups considered taxonomically well studied, awaiting discovery from the region. Further, species like *P. rupestris* sp. nov. which have specific habitat requirements, rills flowing over rocky cliffs in tropical jungle in its particular case, are most vulnerable to climate change. In light of the global insect decline, it behoves the odonatologists of the region to describe species hitherto unknown to science, document their ecology and work towards their conservation expeditiously.

## Acknowledgements

We convey our sincere thanks to the Kerala Forests & Wildlife Department, Government of Kerala, for permitting us to conduct this study (Collection permits Ref: KFDHQ-5492/2021-CWW/WL10 dt 13.04.2022 and KFDHQ-4231/2021-CWW/WL-10 dt 19.12.2022) and providing us with all logistic support. We thank Sujith M, Muneer PK and Madhavan M for assisting us in the field. We thank the Principal, Christ College (Autonomous), Irinjalakuda, for facilitating the laboratory work. We gratefully acknowledge the help of Niranjana KS for the illustrations. The first, second and third authors are grateful to Society for Odonate Studies (SOS) for the encouragement to study Odonata. The first author acknowledges the financial support provided by IUCN Save Our Species (Fondation Segré Conservation Action Fund-Project code: 2022C-2) for conducting fieldwork in Wayanad.

